# The Impact of Colistin Resistance on the Activation of Innate Immunity by Lipopolysaccharide Modification

**DOI:** 10.1101/2022.11.17.517013

**Authors:** José Avendaño-Ortiz, Manuel Ponce-Alonso, Emilio Llanos-González, Hugo Barragán-Prada, Luna Ballestero, Roberto Lozano-Rodríguez, Francesc J. Márquez-Garrido, José María Hernández-Pérez, María-Isabel Morosini, Rafael Cantón, Rosa del Campo, Eduardo López-Collazo

## Abstract

Colistin resistance is caused by different lipopolysaccharide (LPS) modifications, and we propose to evaluate the effect on the innate immune response of *in vivo* and *in vitro* colistin resistance acquisition. We used 2 pairs of isogenic strains: (1) *Escherichia coli* ATCC25922, susceptible to colistin and its isogenic transconjugant-carrying *mcr*-1 gene; and (2) OXA-48, CTX-M-15 *K. pneumoniae* susceptible to colistin (CS-Kp) isolated from a urinary infection and its colistin-resistant variant (CR-Kp) from the same patient after prolonged treatment with colistin. No mutation of described genes for colistin resistance (*pmrA, pmrB, mgrB. phoP/Q* and *crrAB*) were found in the CR-Kp genome; however, LPS modifications were characterized by negative-ion MALDI-TOF. The strains were co-cultured with human monocytes to determine their survival after phagocytosis and induction to apoptosis. Also, monocytes were stimulated with bacterial LPS to study cytokine and immunecheckpoint production. The addition of 4-amino-4-deoxy-l-arabinose (Ara4N) to lipid A of CR-Kp accounted for the colistin resistance. CR-Kp survived significantly longer inside human monocytes after being phagocytosed compared with the CS-Kp strain, whereas no significant differences were observed for the *E. coli* isogenic strains. In addition, LPS from CR-Kp induced both higher apoptosis in monocytes and higher levels of cytokine and immune checkpoint production than LPS from CS-Kp. This effect was strictly the opposite for *E. coli*. Our data reveal a variable impact of colistin resistance on the innate immune system, depending on the responsible mechanism. Adding Ara4N to LPS increases bacterial survival after phagocytosis and elicits a higher inflammatory response than its colistin-susceptible counterpart.

## INTRODUCTION

Lipopolysaccharide (LPS) is a glycolipid from the outer membrane of Gram-negative bacteria that comprise a lipid A domain (endotoxin), a core oligosaccharide, and a repetitive glycan polymer (O-antigen) that projects above the cell surface (1). LPS is the target for colistin (polymyxin E) (2), an old antibiotic that is currently often the only therapeutic option for carbapenemase-producing and antibiotic multidrug-resistant pathogens (3, 4). However, the colistin resistance rate is increasing worldwide (5, 6), mainly as a consequence of LPS modifications following chromosomal mutations in the *phoP/phoQ, pmrA/pmrB*, and *mgrB* genes, or by horizontal acquisition of a plasmid carrying the *mcr*-1 gene (7). Despite the structural modifications, a significant fitness of both chromosomal- (8, 9), and plasmidic-borne (10, 11) mutations leading LPS variants has been reported in several bacterial species, explaining the worldwide expansion of colistin-resistant strains.

LPS is one of the major pathogen-associated molecular patterns involved in the activation of the host innate immune system in the early stages of infection (12–15), and its alterations dampen immune recognition (16, 17). After LPS binds to the TLR-4 receptor, monocytes and macrophages activate an inflammatory response through the NF-κB pathway leading to cytokine production (14, 18). Curiously, LPS also triggers the expression of some immune checkpoints and other factors, leading to an immunosuppressive status known as endotoxin tolerance (19, 20). Several reports point to immune checkpoints as both therapeutic targets and outcome biomarkers in cancer and infectious diseases (21, 22). However, their modulation after LPS exposure has scarcely been explored, particularly for LPS variants of colistin-resistant microorganisms (15). In this regard, most available data are contradictory as some authors considered LPS modification beneficial for bacterial-host interactions and others defined them as detrimental (10, 23–27). Herein, we aim to assess the immunological impact of colistin resistance using pairs of isogenic strains (colistin-susceptible and -resistant) of *Escherichia coli* and *Klebsiella pneumoniae* with different resistance mechanisms.

## RESULTS

### Lipid A modification in CR-Kp

Structural variations of CR-Kp lipid A with respect to that from CS-Kp were determined by negative MALDI-TOF. CS-Kp exhibited the 2 major ions of m/z 1814 and 1841 (Figure 1A), corresponding to the hexaacyl species of lipid A, with a C’-2 acyl-oxo-acyl chain A hydroxylation (-OH), as previously described (28–32). CR-Kp lipid A exhibited an increase of relative intensity of the m/z 1814 peak compared with the CS-Kp. Moreover, CR-Kp had a further ion of m/z 1971 (Figure 1B), resulting from the addition of 4-amino-4-deoxy-l-arabinose (Ara4N) by glycosylation in the C’-1 phosphate group of the m/z 1841 structure with a PRR of 0.29. This latter ion was described as one of the major causes of colistin resistance in *K. pneumoniae* (32–34).

**Figure 1.**
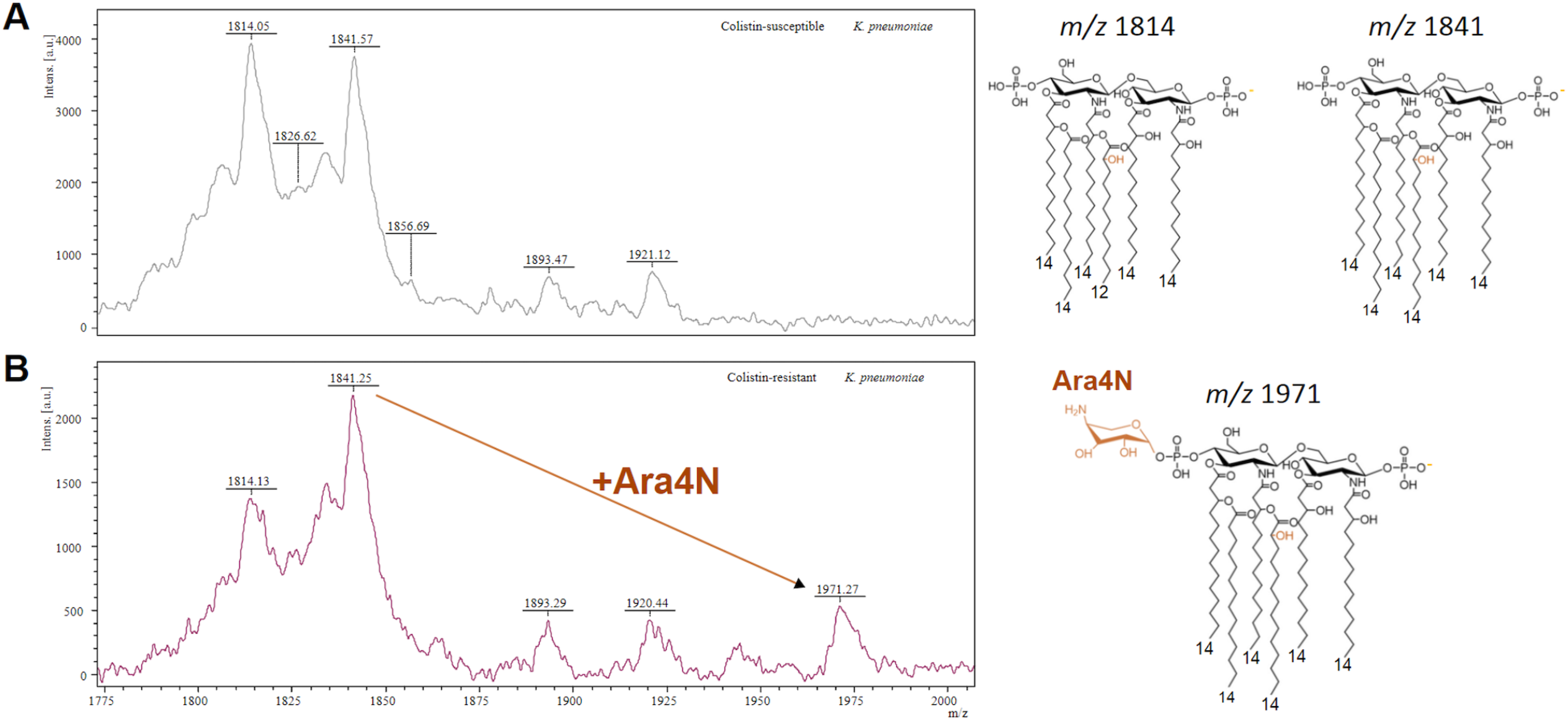
Differential lipid A composition of CS-Kp and CR-Kp strains. Representative negative-ion MALDI-TOF mass spectra of lipid A from the colistin-susceptible **(A)** and colistin-resistant **(B)** *K. pneumoniae* clinical strains. Proposed chemical structures are shown of the most relevant lipid A ions, including the 4-amino-4-deoxy-l-arabinose (Ara4N) addition, explaining their colistin resistance.

### CR-Kp evades host intracellular killing and enhances apoptosis

Although the rates of phagocytosis (Figure 2A) were similar, CR-Kp exhibited a significant intracellular killing evasion, staying alive longer inside the monocytes than CS-Kp (Figure 2B). The control *E. coli* ATCC 25922 strain and its resistant variant by conjugation of the *mcr*-1 plasmid showed no significant difference between them for survival to phagocytosis, and both decreased faster than CR-Kp (Figure 2C). CR-Kp induced significant monocyte apoptosis (Figure 3), particularly total apoptosis, by major survival inside the cells, whereas no differences were found between the *E. coli* strains.

**Figure 2.**
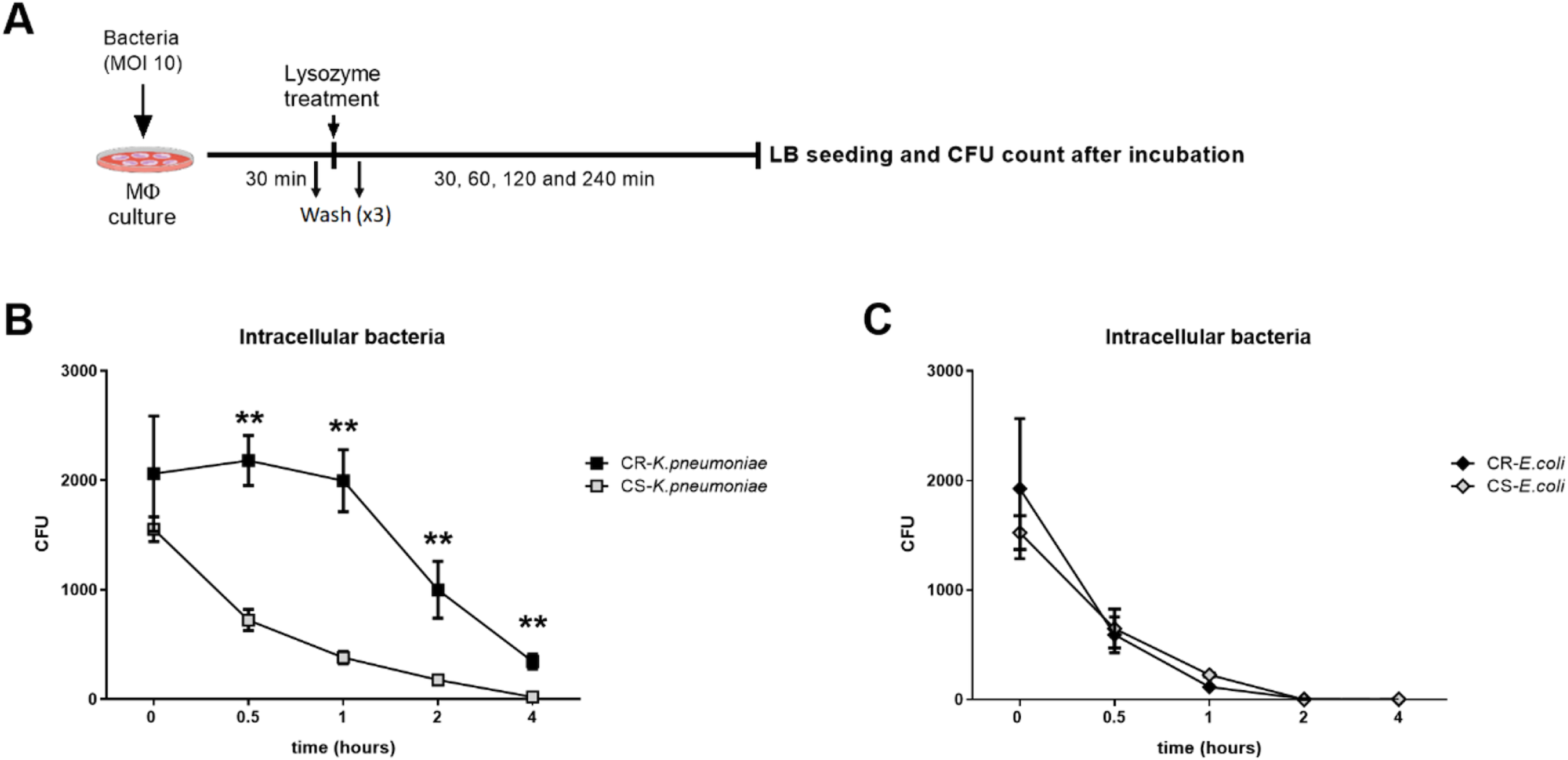
The CR-Kp strain exhibited higher intracellular killing evasion ability in a phagocytosis assay. **(A)** Schematic representation of procedures for phagocytosis and intracellular killing assays. Human monocytes were coinfected with separate strains at multiplicity of infection of 10 (10 bacteria per monocyte) for 30 min. Then, the culture was disrupted with lysozyme and washed to discard non-phagocytized bacteria. Total colony forming units (CFUs) of *K. pneumoniae* **(B)** and *E. coli* **(C)** inside the monocyte at various time points after phagocytosis. **, *p*<0.01 in Mann-Whitney U test for CR and CS comparison, n=4 independent biological replicates with two technical replicates each.

**Figure 3.**
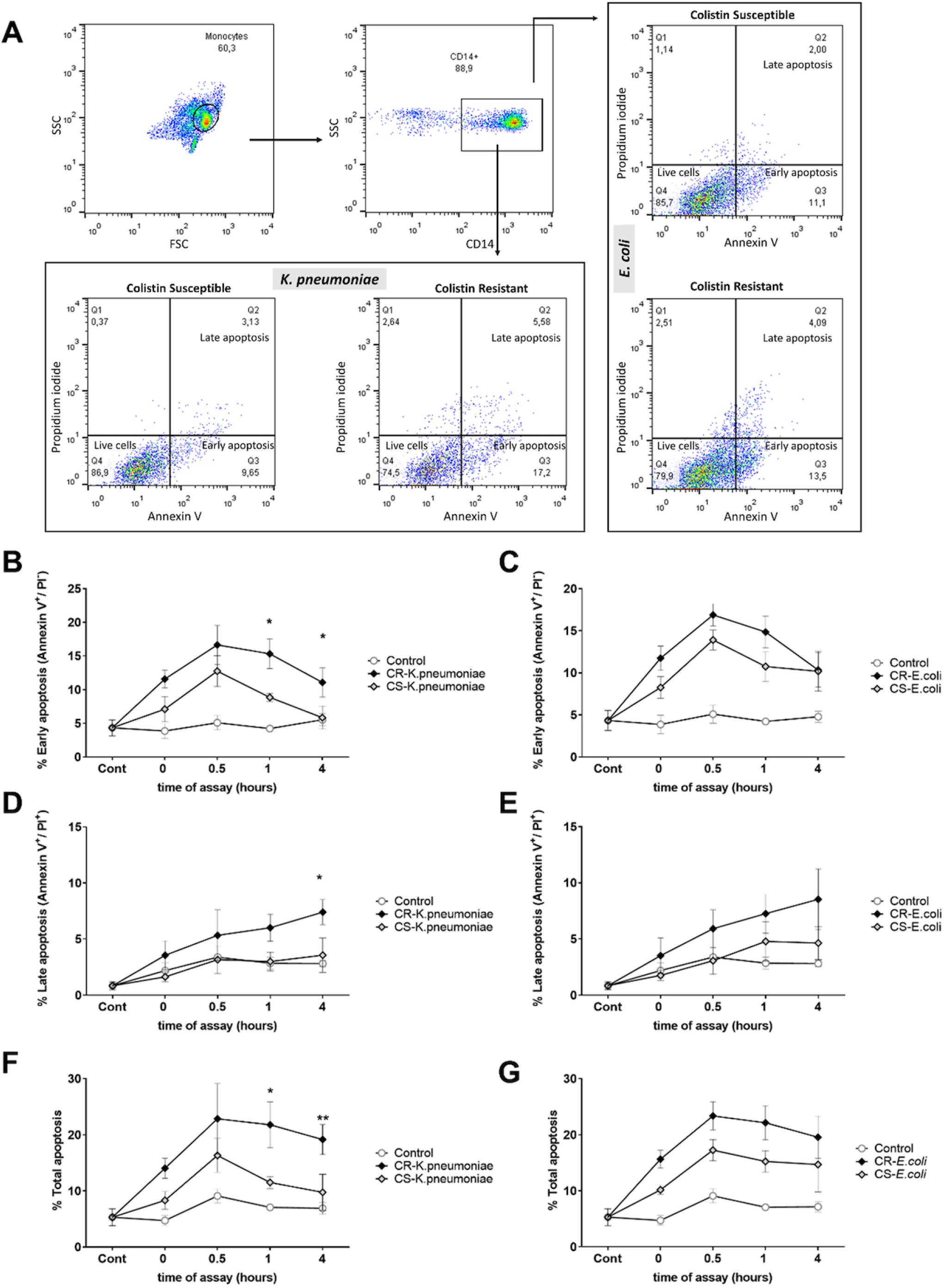
The CR-Kp strain induced higher apoptosis in human monocytes. **(A)** Representative gating strategy followed in the apoptosis assay. Percentage of early **(B** and **C**),late **(D** and **E**), and total **(F** and **G**) apoptosis in gated CD14^+^ monocytes at various time points from coculture at a multiplicity of infection of 10 (10 bacteria per monocyte) with colistin resistant (CR) and colistin susceptible (CS) *K. pneumoniae* and *E. coli*. **, *p*<0.01 in Mann-Whitney U test for CR and CS comparison, n=4 independent biological replicates.

### CR-Kp LPS triggered increased cytokine production and immune checkpoints, contrary to what was observed in CR-Ec LPS

We next assessed whether the lipid A modified LPS elicited a different immune response. We found LPS from CR-Ec induced a lower inflammatory response by lower production of tumor necrosis factor (TNF)α, interleukin (IL)-6, and C-X-C motif chemokine ligand (CXCL)10 (Figure 4) compared with CS-Ec in human monocytes. In contrast, for *K. pneumoniae*, an increment of TNFα, IL-6, and CXCL10 production by LPS stimulation with LPS from the CR-Kp variant was observed (Figure 4). Other cytokines, such as IL-1 beta, IL-10, and interferon-gamma, showed the same pattern in both *E. coli* and *K. pneumoniae*, but without statistical significance (Supplementary Figure 1).

**Figure 4.**
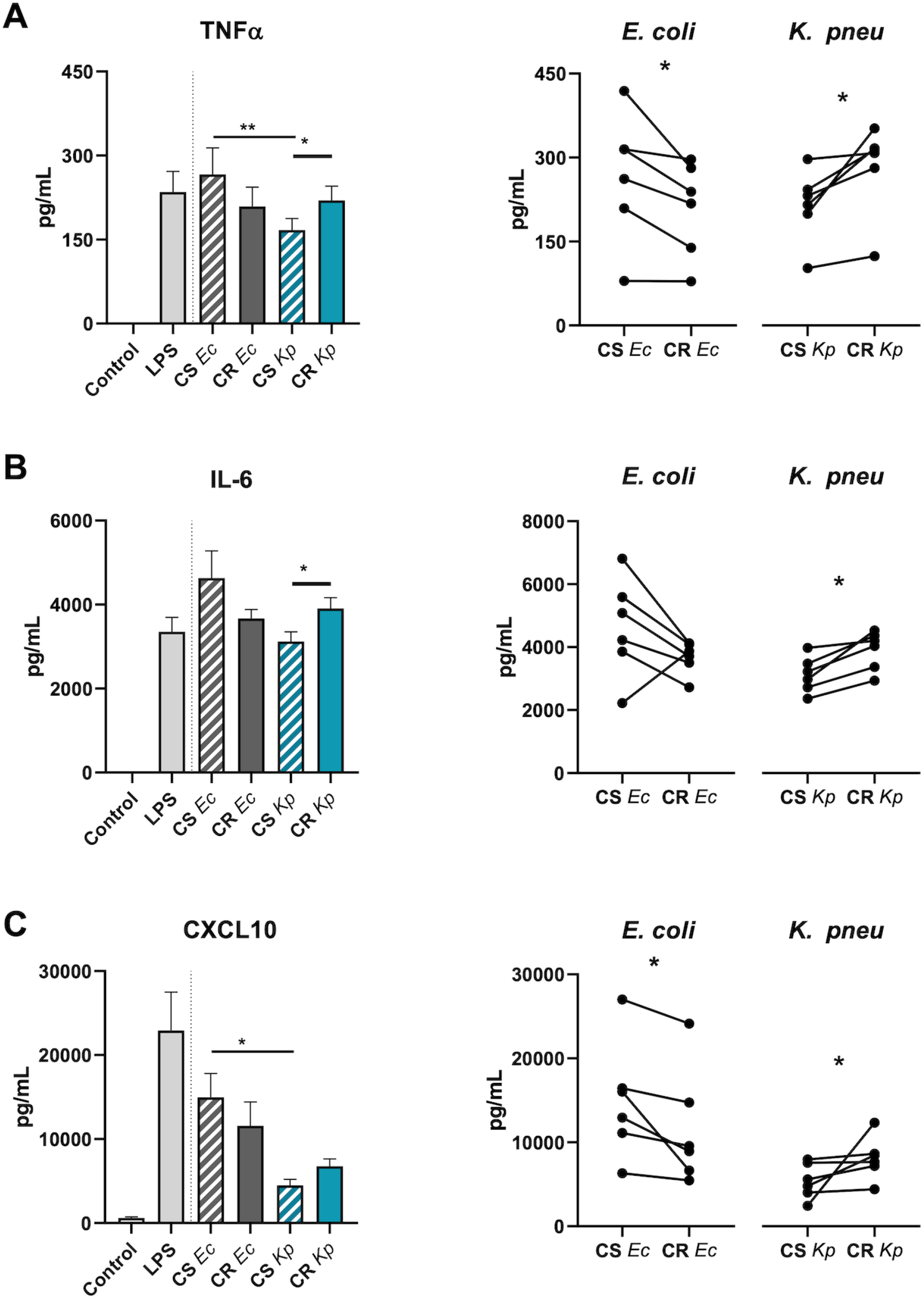
LPS from CR-Kp induced higher inflammatory cytokine production. Levels of TNFα **(A)**, IL-6 **(B)**, and CXCL10 **(C)** in monocyte supernatant after 24 h of stimulation with 10 ng/mL of commercial LPS (LPS, from *E. coli* Olll:B4) or the same concentration of isolated LPS from all 4 strains. *, *p*<0.05 and **, *p*<0.01 in Friedman test followed by Dunn for multiple group comparison and Wilcoxon test for paired CS and CR comparison, n=6 independent biological replicates.

Several reports have indicated the potential role of immune checkpoints in the context of sepsis, given that LPS can induce their expression. We studied the changes in monocyte supernatant after stimulation with the purified LPS of our strains. A significantly higher overall production of galectin-9, sCD25, sTim-3, and sCD86 were observed for CR-Kp LPS compared with that from the susceptible variant. In *E. coli*, the LPS of CR-Ec induced only lower production for sTim-3 with respect to CS-Ec (Figure 5, Supplementary Figure 2).

**Figure 5.**
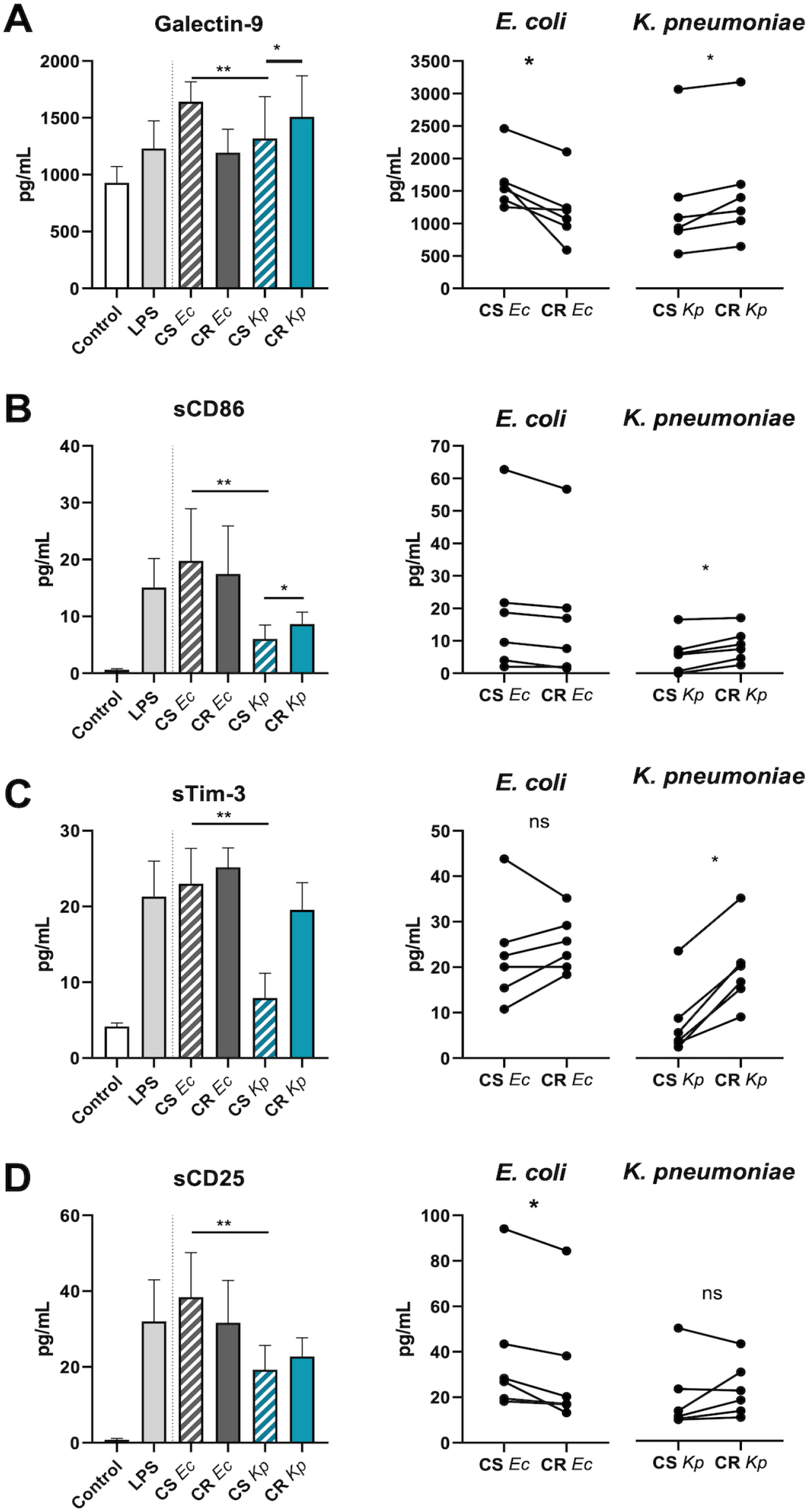
LPS from CR-Kp strain induced higher immune checkpoint production. Levels of galectin-9 **(A)**, soluble CD25 **(B)**, soluble Tim-3 **(C)**, and soluble CD86 **(D)** in monocyte supernatant after 24 h of stimulation with 10 ng/mL of commercial LPS (LPS, from *E. coli* O111:B4) or the same concentration of isolated LPS from all 4 strains. *, *p*<0.05; **, *p*<0.01 in Friedman test followed by Dunn for multiple group comparison and Wilcoxon test fosr paired CS and CR comparison, n=6 independent biological replicates.

## DISCUSSION

The aim of this study was to decipher the impact of colistin resistance on the innate immune system caused by lipid A modification from isogenic pairs of *E. coli* and *K. pneumoniae* strains and their purified LPS in monocytes. Previous studies had revealed that the *mcr*-1 gene harboring bacteria induced less inflammation than the wild type (10, 35) although its effect on bacterial phagocytosis was not defined. A more recent study focusing on the *mcr-3* gene found that their expression conferred protection against phagocytosis by macrophages. Here, we did not find differences in bacterial phagocytosis or intracellular killing in *mcr*-1 carrying *E. coli* compared with its wild-type isogenic strain.

In contrast, the CR-Kp strain exhibited clear resistance against intracellular killing that was considerably higher than its susceptible counterpart, although it also induced phagocytosis at a higher rate than its susceptible counterpart. The observed difference in phagocytosis between CR-Ec and CR-Kp could be explained by their lipid A modifications. Negative-ionization MALDI-TOF revealed that CR-Kp exhibited an addition of Ara4N to hydroxylated lipid A, whereas in *E. coli, mcr*-1 expression has been shown to cause a phosphoethanolamine addition to lipid A (36). The PRR of our CR-Kp was 0.29, a similar value to the observed in Ara4N lipid A modification from colistin-resistant OXA-48-producing *E. coli* that lacked *mcr* genes (37).

Resistance to colistin is associated with LPS modifications altering the immunogenic nature of lipid A as a result of enzyme synthesis mutations (10, 15, 27, 38). It is important to note that we did not find plasmid genes, nor mutations on the *pmrB, pmrA*, or *mgrB* genes, nor other reported mechanisms, including *pho*P/Q mutations in CR-Kp (39); thus, the subjacent genotype for this modification remains elusive. Dogan *et al*. described that the iron uptake systems by *kfu* and *ybtS* genes in colistin-resistant *K. pneumoniae* clones enhanced survival against the phagocytosis of neutrophils (40). Similarly, the absence of WcaJ glycosyltransferase in *K. pneumoniae* with the subsequent reduction of colanic acid in LPS reduced phagocytosis ability. Along these lines, although we performed a paired study with 2 pairs of isogenic strains, a possible study limitation is that we did not include all the possible mechanisms for colistin resistance. The effect on the immune response in other forms of resistance acquisition should be evaluated in future studies.

The LPS molecule binds the TLR-4 receptor in innate immune cells, triggering the inflammatory response through cytokine production via the NF-κB pathway. Infection with recombinant *mcr*-1-expressing *E. coli* significantly downmodulated p38-MAPK activation and NF-κB nuclear translocation, leading to a decrease in the proinflammatory cytokines TNF-α, IL-12, and IL-1β, and lower caspase 1 activity with respect to the colistin-susceptible strains (32). In the same manner, LPS from *E. coli mcr*-1 induced lower production of IL-6 and TNF compared with parental LPS (10). However, these CR mutants showed reduced fitness and attenuated virulence in a *Galleria mellonella* model (10). We used *mcr*-1 *E. coli* as a control, as well as to reduce inflammatory cytokine production; however, no patent differences were found for monocyte apoptosis induction. In contrast, our *K. pneumoniae* strains had a completely opposite effect characterized by higher inflammatory cytokines as well as higher apoptosis induction in the CR variant. Our *K. pneumoniae* results therefore diverge from those observed in *E. coli* and are closer to those observed in *Acinetobacter baumannii*, in which multiresistant mutants induced more inflammation and resisted phagocytosis more than wild-type variants (35). Moreover, modifications of lipid A in the 2 bacterial species could be different. The 4-AraN addition in our clinical *K. pneumoniae* strain appears to be responsible for colistin resistance. This same modification in *Burkholderia* lipid A is described to enhance inflammatory responses in macrophages (36,37), similarly to what we found in *K. pneumoniae*.

Immune checkpoints either activate or inhibit the immune system (22, 38); their blockade is a successful treatment for lung and melanoma cancer (38, 39). Although bacterial LPS had been shown to also induce the expression of immuno-checkpoints (19, 20, 40) the effects of LPS modifications remained unexplored, particularly those related to colistin-resistance. Our study demonstrated that colistin resistance through *in vivo* LPS modification induced higher sCD25, sCD86, galectin-9, and sTim-3 production, probably causing detrimental effects in the host immune response; i.e., macrophagic syndrome is defined by high levels of sCD25 (41). For its part, galectin-9 and sTim3 could play a role in immune system anergy, according to data indicating that these receptors are linked to secondary infection, septic shock, and outcomes in infectious diseases (42–44). Therefore, the relationship between colistin resistance, immune checkpoint blockade, and macrophage activation syndrome in infectious diseases should be explored.

In conclusion, despite their isogenic background, a different immune activation profile in terms of phagocytosis, intracellular killing evasion, inflammatory response, and immune checkpoint expression were demonstrated for colistin resistant *E. coli* and *K. pneumoniae* strains. In addition, colistin resistance in *K. pneumoniae* significantly delayed internal bacteria clearance by human monocytes, and their LPS induced a higher inflammatory response and immune-checkpoint expression. Our data described the immunological cost of colistin resistance by our specific LPS modification, although these effects should be individually studied, taking account each type of LPS modification in the various bacterial species.

## MATERIAL AND METHODS

### Bacterial strains

The 2 sets of isogenic bacterial strains and their minimal inhibitory concentration colistin values consisted of an *E. coli* colistin-susceptible ATCC 25922 strain (CS-Ec, 0.5 mg/L) and its resistant variant by *mcr-1* plasmid conjugation (CR-Ec, 4 mg/L); and *K. pneumoniae* susceptible (CS-Kp, 0.5 mg/L) and resistant (CR-Kp, 32 mg/L) clinical strains. The CS-Kp was a clinical OXA-48 and CTX-M-15-producing *K. pneumoniae* strain obtained from the urine of a 79-year-old hospitalized woman with diabetes mellitus and stage IV chronic kidney disease who had recurrent urinary tract infections. Although the patient was successfully treated with meropenem plus amikacin plus colistin, she relapsed 2 months later with a colistin-resistant variant, the CR-Kp isolate. The CS-Kp and CR-Kp were submitted to whole-genome sequencing on an Illumina MiSeq platform (2×300 bp) at mean read depth of 158x. Briefly, genome was assembled with VelvetOptimiser 2.2.5 and SPAdes v3.9.0. Prokka 1.12-beta to annotate susceptible *K. pneumoniae* assembled genome, and BWA v0.7.5 and Samtools v0.1.18 for mapping CR-Kp against susceptible reference genome. Variants were analyzed by VarScan with no mutations detection of the previously reported genes *pmrA, pmrB*, *mgrB. phoP/Q* and *crrAB* responsible for colistin resistance. Fitness cost of the colistin resistance acquisition in *K. pneumoniae* strains were tested in a previous work (32) finding a relative growth rate of 0.973 and a fitness decrease of 2.66% (*p* value 0.059) in CR-Kp compared with CS-Kp.

### Mass spectrometry characterization of *K. pneumoniae* lipid A

Lipid A from the *K. pneumoniae* clinical strains was extracted with the MBT Lipid Xtract kit (Bruker Daltonics, Germany), following the manufacturer’s instructions. Briefly, bacteria were grown overnight in Mueller-Hinton agar, picked up by a 1-μL inoculation loop, and mixed in 50 μL of MTB Lipid Xtract Hydrolysis buffer in a low-binding tube. Then, 44 μL of the cell suspension was discarded, and the remaining 6 μL were submitted to a heating process at 90 °C for 10 min. The tubes were left for 2 min with the lid open to completely evaporate the buffer. The dried pellets were washed with 50 μL of washing buffer without dissolving the pellet. The total volume of the washing buffer was discarded by pipetting. Finally, 5 μL of matrix was pipetted up and down for 15-20 seconds to resuspend the dried pellet, and 2 μL were spotted on an MTP 364 polished steel MALDI target plate. Mass spectrometry runs were performed with a matrix-assisted laser desorption/ionization time-of-flight (MALDI-TOF) autoflex maX spectrometer (Bruker Daltonics) in linear negative-ion mode. Spectra were analyzed using FlexAnalysis v.3.4. software (Bruker Daltonics). The polymyxin resistance ratio (PRR) were estimated from the acquired spectra by summing the intensities of the lipid A peaks attributable to the addition Ara4N (m/z 1971) and dividing this number by the intensity of the peak corresponding to native lipid A (m/z 1814) as described by Furnis *et al*. (30).

### LPS isolation

The LPS was purified following a modified version of a previously described method (44). Briefly, 1.5 mL of overnight cultures at 37 °C in Luria–Bertani (LB) broth (Difco, USA) of both the susceptible and resistant strains (supplementing with 6 mg/L of colistin) adjusted to an OD_600_=0.5 were centrifuged at 10,600 x *g* for 10 min. The pellet was suspended in 200 μl of sodium dodecyl sulfate (SDS) buffer dyed with bromophenol (a solution of 4%β-mercaptoethanol, 4% SDS, and 20% glycerol in 0.1 M Tris-HCl, pH 6.8, diluted 1:1 in sterile distilled water), and then boiled for 15 min. After room tempering, 10 μl of a proteinase K solution (10 mg/L) was added, incubating it at 59 °C for 3 h. Afterward, 200 μl of ice-cold tris-saturated phenol was added to each sample, with vigorous vortexing. Then, samples were incubated at 65 °C for 15 min, shaking occasionally. Again, at room temperature 1 mL of diethyl ether was added to each sample, which was then vortexed and centrifuged at 16,000 x *g* for 10 min. Finally, the bottom blue layer containing the LPS was extracted. The product was extracted once again, starting from the SDS step, to obtain a purified solution of LPS.

Purity was assessed by SDS gel and coomassie blue staining as previously described (45). LPS concentration was determined by a quantitative chromogenic endpoint test, the QCL-1000 Assay (Lonza Walkersville, Inc., Walkersville, MD, USA).

### Ethics Statement

The Ethics Committee of Ramón y Cajal University Hospital approved the epidemiological study of the multiresistant antibiotic isolates. The human blood buffy coats from healthy anonymous donors were obtained from the blood donor Service of La Paz University Hospital. All the participants provided written consent in accordance with the ethical guidelines of the 1975 Declaration of Helsinki and the Committee for Human Subjects of La Paz University Hospital (PI-4100).

### Peripheral blood monocyte isolation

Healthy donors were recruited from the blood donor service of La Paz University Hospital, and fresh blood from venipuncture was collected in K2EDTA tubes (BD Vacutainer, USA). The blood was added to Ficoll-Paque PLUS (Cytiva, MA, USA) according to the manufacturers’ protocol. The peripheral blood mononuclear cells were counted, and the monocyte percentage was checked by flow cytometry (BD FACSCalibur). The monocytes population were enriched by an adherence protocol after 1 h of culture in serum-free Roswell Park Memorial Institute (RPMI) liquid medium as previously described (20,46).

### Phagocytosis and host intracellular killing assay

Phagocytosis was performed following previously described reports (46, 47). The schematic procedure is shown in Figure 2A. Briefly, isolated human monocytes were exposed to the bacteria at a multiplicity of infection (MOI) of 10 (10 bacteria per monocyte) in RPMI without antibiotics for 30 min. The monocytes were washed three times with PBS and treated with lysozyme (1 mg/mL) for 10 min and washed another three times to remove the non-phagocyted bacteria. Supernatant after these washes were seeded on LB agar plates (Difco) at 37 °C for 24 h as negative controls. To determine the intracellular bacteria, 0, 1, 2, and 4 h after phagocytosis, monocytes were washed twice and subjected to soft lysing with 0.5% Triton for 10 min. Aliquots from monocytes lysate were seeded on LB agar plates (Difco) at 37 °C for 24 h for counting colony forming units.

### Flow cytometry staining and apoptosis assay

Monocytes resulting from the host intracellular killing assay described above were used to analyze apoptosis induction.

Briefly, monocytes were washed twice with phosphate-buffered saline (PBS) after lysozyme treatment and cultivated for 1 and 4 h. Then, cells were collected and labeled with CD14 APC (Immunostep, Spain). Apoptosis was determined employing the fluorescein isothiocyanate Annexin V Apoptosis Detection Kit (Immunostep) following the manufacturer’s protocol. Additionally, an apoptosis assay was performed on monocytes from healthy donors by stimulation for 1 and 4 h with 100 ng/mL of LPS purified from the bacterial strains.

### Cytokine and immune checkpoint quantification

The supernatants of the monocyte cultures were analyzed after stimulation with 10 ng/mL of the different purified LPSs for 24 h. Commercial *E. coli* O111:B4 LPS (Sigma-Aldrich, MA, USA) was used as a positive control. The inflammatory cytokines and the soluble immune checkpoints production in supernatant were quantified by a multiplex cytometric bead array (LEGENDplex HU Essential Immune Response Panel and LEGENDplex HU Immune Checkpoint Panel 1, respectively). Samples were acquired in a BD FACSCalibur cytometer and analyzed by the LEGENDplex Data Analysis Software Suite (Qognit, Inc., CA, USA).

### Statistical analysis

The results are expressed as mean ± standard error of the mean. The sample size (n) as well as the statistical test for each experiment are shown in the Figure legends. All experiments were repeated at least twice, in different replicas and using monocytes from at least 4 unrelated donors. Briefly, for the phagocytosis experiments, data were first compared using the Kruskal–Wallis test, followed by the Mann–Whitney U test. For cytokine and immune checkpoint production after stimulation with purified LPS from the CS and CR variants, a Friedman test and Dunn’s multiple comparison test, as well as a Wilcoxon signed-rank test, were performed. *P*-values of <0.05 were considered significant at a 95% confidence interval. Statistical analyses were performed with IBM SPSS 23 (Armonk, NY, USA) and GraphPad Prism 8 (San Diego, CA, USA) software.

## ACKNOWLEDGEMENTS

We are indebted to the transfusion center of the Community of Madrid, and grateful to Marta Cobo for excellent technical assistance. The support of Santos del Campo is also acknowledged.

## FUNDING

This study was supported by Fundación La Paz-HULP and by the Instituto de Salud Carlos III (ISCIII), PI20/00164 to RdC, co-funded by the European Union. JAO is supported by a Sara Borrell contract (CD21/00059) and MPA by a Rio Hortega contract (CM19/00069), both from the Instituto de Salud Carlos III.

